# Molecular epidemiology and drug resistance patterns of *Mycobacterium tuberculosis* complex isolates from university students and the local community in Eastern Ethiopia

**DOI:** 10.1101/322958

**Authors:** Abiyu Mekonnen, Matthias Merker, Jeffrey M Collins, Desalegn Addise, Abraham Aseffa, Beyene Petros, Gobena Ameni, Stefan Niemann

## Abstract

**Background:** Previous studies suggest the burden of pulmonary tuberculosis (PTB) in Ethiopia may be greater in university students relative to the overall population. However, little is known about the transmission dynamics of PTB among students and members of the communities surrounding university campuses in Eastern Ethiopia.

**Methods:** A cross sectional study was conducted in Eastern Ethiopia among culture-confirmed PTB cases from university students (n=36) and community members diagnosed at one of four hospitals (n=152) serving the surrounding area. Drug susceptibility testing (DST) was performed on Mycobacterium Tuberculosis Complex (MTBC) isolates using BD Bactec MGIT 960 and molecular genotyping was performed using spoligotyping and 24-loci MIRU-VNTR. MTBC strains with Identical genotyping patterns were assigned to molecular clusters as surrogate marker for recent transmission and further contact tracing was initiated among clustered patients.

**Results:** Among all study participants, four MTBC lineages and 11 sub-lineages were identified, with Ethiopia_3 being most common sub-lineage (29.4%) and associated with strain clustering (P= 0.016). We identified 13 (8.1%) strains phylogenetically related to the known Ethiopian sub-lineages with a distinct Spoligotyping patterns and designated as Ethiopia_4. The clustering rate of MTB strains was 52.9% for university students and 66.7% for community members with a Recent Transmission Index (RTI) of 17.6% and 48.4%, respectively. Female gender, urban residence, and new TB cases were significantly associated with strain clustering (p<0.05). Forty-eight (30%) of the study participants were resistant to one or more first line anti TB drugs, three patients were classified as multidrug resistant (MDR), defined by isoniazid and rifampicin resistance.

**Conclusion:** We found evidence of significant PTB cases clustering and recent transmission among Ethiopian university students and the local community in eastern Ethiopia; with Ethiopia_3 being the predominant circulating sub-lineage. A country wide comprehensive molecular surveillance and drug resistance profiling of MTBC strains and Implementation of TB control programs within universities and the surrounding community should be considered to decrease TB transmission.

## Background

Tuberculosis (TB) remains a major threat to public health worldwide [1], with an estimated 10.4 million cases in 2016 [2] Ethiopia is one of the fourteen countries with the highest TB burden with an annual incidence rate of 177/100,000 population [2] Young adults in Ethiopia have a higher TB incidence than any other age group [3], which has a negative impact on the country’s socio-economic development [4]. Halting TB transmission and decreasing TB incidence rates, especially in young adults, will be an essential component of the public health program in Ethiopia. However, in communities of young adults, such as university students, little is known about recent PTB infection rates and the primary drivers of TB transmission. These communities offer an opportunity for interrupt transmission with infection control measures such as TB screening and contact tracing.

Molecular strain typing (genotyping), namely 24-loci Mycobacterial Interspersed Repetitive Units-Variable Number Tandem Repeat (MIRU-VNTR) technique in combination with Spacer Oligo Nucleotide typing (Spoligotyping), is widely used to investigate local transmission dynamics and evaluate TB control programs [5,6,7]. In addition, certain MTBC lineages (e.g. lineage 2 [Beijing]) have been associated with increased pathogenicity and resistance to specific drugs, which increase the risk of drug resistant MTBC strains in a region [8].

Recent studies on university students in central Ethiopia suggested TB incidence to be much higher on school campuses relative to surrounding communities [9]. Rapid increases in university enrollment in Ethiopia has led to crowded congregate living environments on college campuses with the potential to facilitate TB transmission. However, the relative contribution of recent TB transmission, both on campus and in the surrounding community, to active TB disease among Ethiopian university students is unknown. We studied university students and surrounding community members diagnosed with pulmonary TB at three universities and four hospitals in eastern Ethiopia to determine the genotypic characteristics, transmission dynamics and drug resistance patterns of MTBC strains circulating in the region.

## Materials and Methods

### Study design and period

A cross-sectional study was conducted among students of three Eastern Ethiopian universities and members of the surrounding community diagnosed with pulmonary tuberculosis (PTB). Inclusion criteria was a positive sputum culture for MTBC. Students with PTB were identified through active case finding between May 2016 and April 2017; community TB cases were enrolled from hospitals serving the geographic areas surrounding the universities from January to April 2017. Participants for whom a valid genotype could not be obtained were excluded from the study. This included evidence of a mixed infection or laboratory cross-contamination as indicated by double alleles at two or more loci during MIRU/VNTR typing and two or more loci with missing data following at least two independent PCR amplifications. Reasons for missing loci included insufficient DNA concentration [10] and nucleotide polymorphisms in the sequence complementary to the PCR primers [11].

### Study sites

The study comprises three regional states and one administration (Oromia, Somali and Harari regional states and Dire Dawa City administration). The towns where the study hospitals were located are: Harar (Harari region), Haramaya (Haramaya district, Oromia region), Dire Dawa (Dire Dawa city administration) and Jigjiga (Somali region). According to the Central Statistical agency of Ethiopian population projection values of 2017, the population of Harari and Somali regional states were 246,000 and 5,748,998, respectively; whereas that of Dire Dawa city administration and Haramaya district were 466,000 and 361,787, respectively [12]. The universities studied were Haramaya University, Dire Dawa University and Jigjiga University.

### Collection of MTBC strains

Students: All full-time students attending Haramaya University, Dire Dawa University and Jigjiga University were screened for PTB by active case finding through dormitory-to-dormitory visits using WHO TB screening document [13] between May 2016 and April 2017. Two spot sputum samples were collected from students with a positive WHO symptom screen. One sputum sample was processed for Acid Fast Bacilli (AFB) sputum smear microscopy and the other one was transported to Harari Health Research and Regional Laboratory, Harar, Ethiopia for MTB culture. Thirty-six student AFB cultures were positive for MTBC and genotype analysis was performed on all isolates.

Local community: Persons diagnosed with PTB from the community surrounding the universities studied were enrolled at four hospitals: Haramaya district hospital (Haramaya), Hiwot Fana specialized university hospital (Harar), Dil Chora hospital (Dire Dawa) and Karamara hospital (Jigjiga). Persons presenting to these facilities with symptoms of PTB and found to have a positive AFB sputum smear between January and April 2017 were approached for study enrollment. Persons giving consent for study participation were administered a standard questionnaire to collect information about relevant clinical and sociodemographic data. An early morning sputum sample was collected from the respective smear positive patient and stored at −20°c until transported to Harari Health Research and Regional Laboratory for TB culture. Initially 171 sputum samples were collected from smear positive PTB cases from the four hospitals, of which 19 (11.3%) samples were excluded due to growth of contaminating flora.

### Laboratory methods

Sputum samples were collected from students with a positive symptom screen and community members with symptoms and a positive AFB sputum. All specimens were cultivated on LJ (BBL™ Lowenstein-Jensen) media at Harari Health Research and Regional laboratory following the standard operational procedures. Isolates on LJ transported to the National TB reference laboratory, Addis Ababa, were reactivated and phenotypic Drug Susceptibility Testing (DST) was performed using the MGIT SIRE kit at a critical concentration of streptomycin (STM) 1 μg, Isoniazid (INH) 0.1 μg, Rifampicin (RIF) 1 μg and Ethambutol (EMB) 5 μg on liquid Mycobacterium Growth Indicator Tube system (MGIT) 960 as described by the manufacturer [14].

DNA from MTBC isolates were extracted and transported to Borstel Molecular Mycobacteriology laboratory, Germany for genotype analysis. Molecular characterization of all isolates was conducted using Spoligotyping [15] and 24- loci MIRU-VNTR customized kits (Genoscreen, Lilli, France) [16]. Spoligotypes common to more than one strain were designated as shared types (ST) and were assigned a shared international type number (SIT) according to the international spoligotype database SpolDB4 [17]. Basic strain classification and MLVA MTBC 15-9 nomenclature assignment was done using the MIRUVNTRplus database [18]. Samples with complete spoligotyping and MIRU/VNTR-24 results were used for clustering analysis. A cluster was defined as two or more MTBC isolates sharing identical 24-loci MIRU/VNTR and spoligotyping patterns. Samples with no PCR amplicon at only one locus in the 24-loci MIRU/VNTR analysis were included for further analysis by considering missing data at the respective locus [19].

## Data analysis

MTBC genotypes were classified in a phylogenetic tree (based on 24-loci MIRU-VNTR profiles) in relation to a MTBC reference collection hosted on the MIRU-VNTRplus website (available at www.miru-vntrplus.org) and considering genotype specific Spoligotyping patterns [18]. Minimum Spanning Trees (MST) were calculated with BioNumerics (Version 7.5; Applied Maths, Sint-Martens-Latem, Belgium) as recommended by the manufacturer (available at http://applied-maths.com). A dendrogram was generated using the Unweighted Pair Group Method with Arithmetic averages (UPGMA) based on the 24-loci MIRU-VNTR profiles. The UPGMA tree was further processed using EvolView, an online visualization and management tool for customized and annotated phylogenetic trees [20] (Fig 1). The Recent Transmission Index (RTI) was calculated as number of clustered patients minus number of clusters divided by total number of patients.

**Fig. 1.**
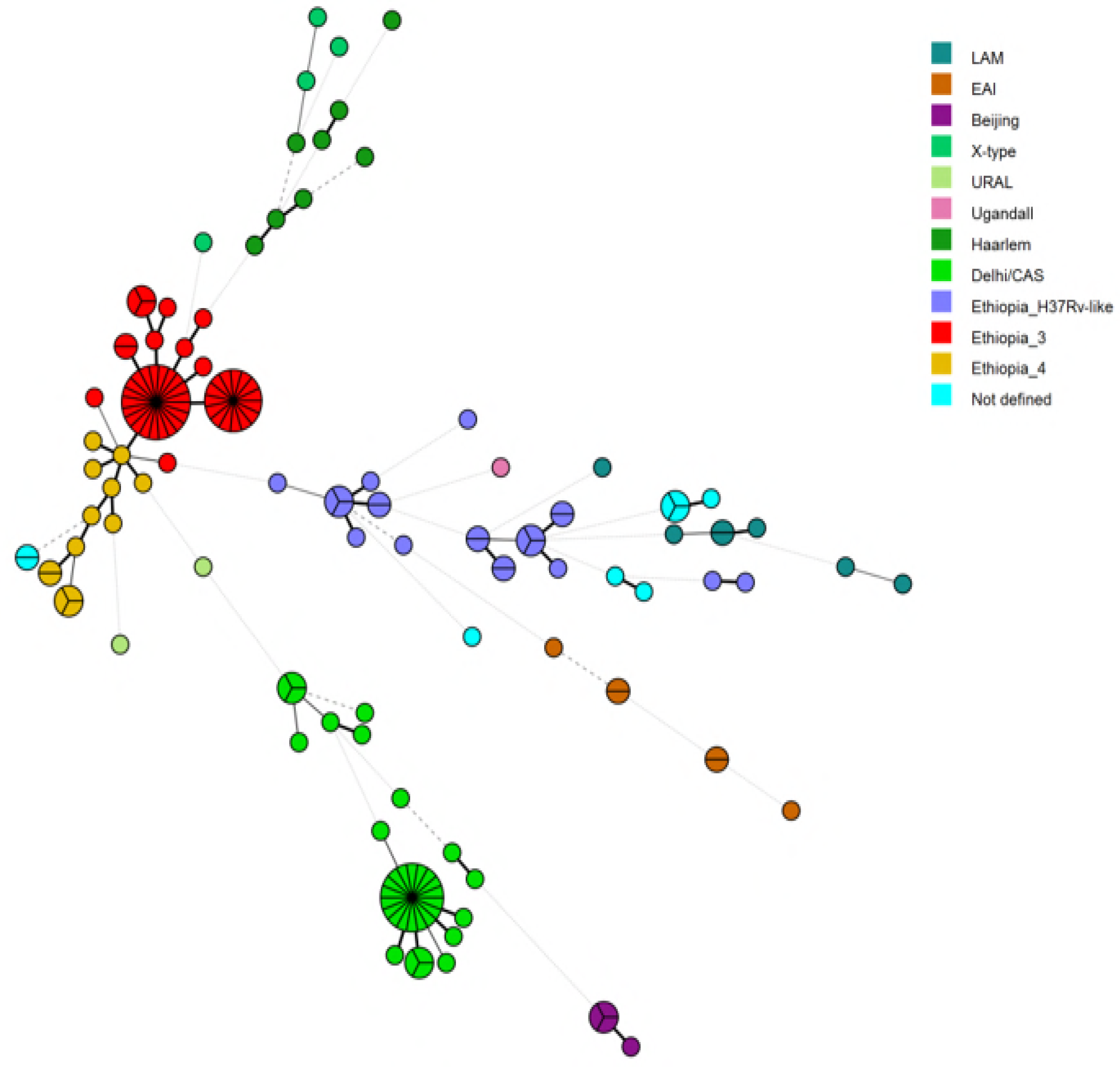
Minimum spanning tree based on MIRU/VNTR profiles of 24- loci of 160 MTBC isolates in eastern Ethiopia. A circle representing a specific genotype is divided in to the number of strains clustering in it. EAI: East Africa India, LAM=M. tuberculosis Latin American Mediterranean.

Data were entered and analyzed using IBM SPSS version 23 statistical package software. Logistic regression was used to estimate the strength of association between strain clustering and different variables. A Chi-square test was used for bivariate analysis of categorical variables. P-values <0.05 were considered as statistically significant. Those factors significantly associated with clustering in the univariate analysis were included in the multivariate regression model.

## Ethical Considerations

The study was ethically reviewed and approved by Addis Ababa University, College of Natural Sciences Research Ethics Review Board. Written informed consent that included information about the risks and benefits of the study was a prerequisite for all university students and local community study participants.

## Results

### Study population

We enrolled 188 study participants with PTB, all of whom were sputum culture positive for MTBC. Molecular genotyping was performed on all isolates using Spoligotyping and MIRU/VNTR-24 loci techniques. Of these, 28 isolates were excluded for technical reasons; two were found to be a mixed infection as indicated by double alleles at two or more loci during MIRU/VNTR typing and 26 isolates had two or more loci missing following PCR amplification. Therefore, 160 MTBC strains were included in the final genotypic analysis: 34 from university students and 126 from the local community. The majority of the study participants were male (80%) and rural residents (60.6%). Forty-six (28.7%) of the participants had previous treatment for TB, and 10.6% were HIV-positive (Table 1).

**Table 1.**
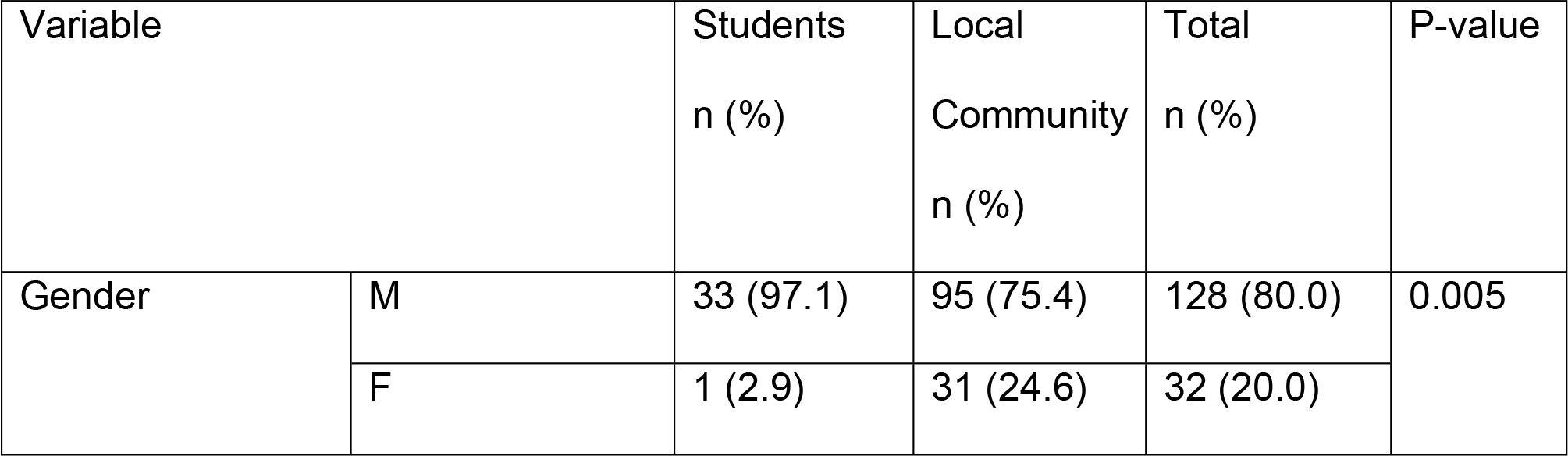
Socio-demographic and other characteristics by study participants, eastern Ethiopia, May 2016 to April 2017

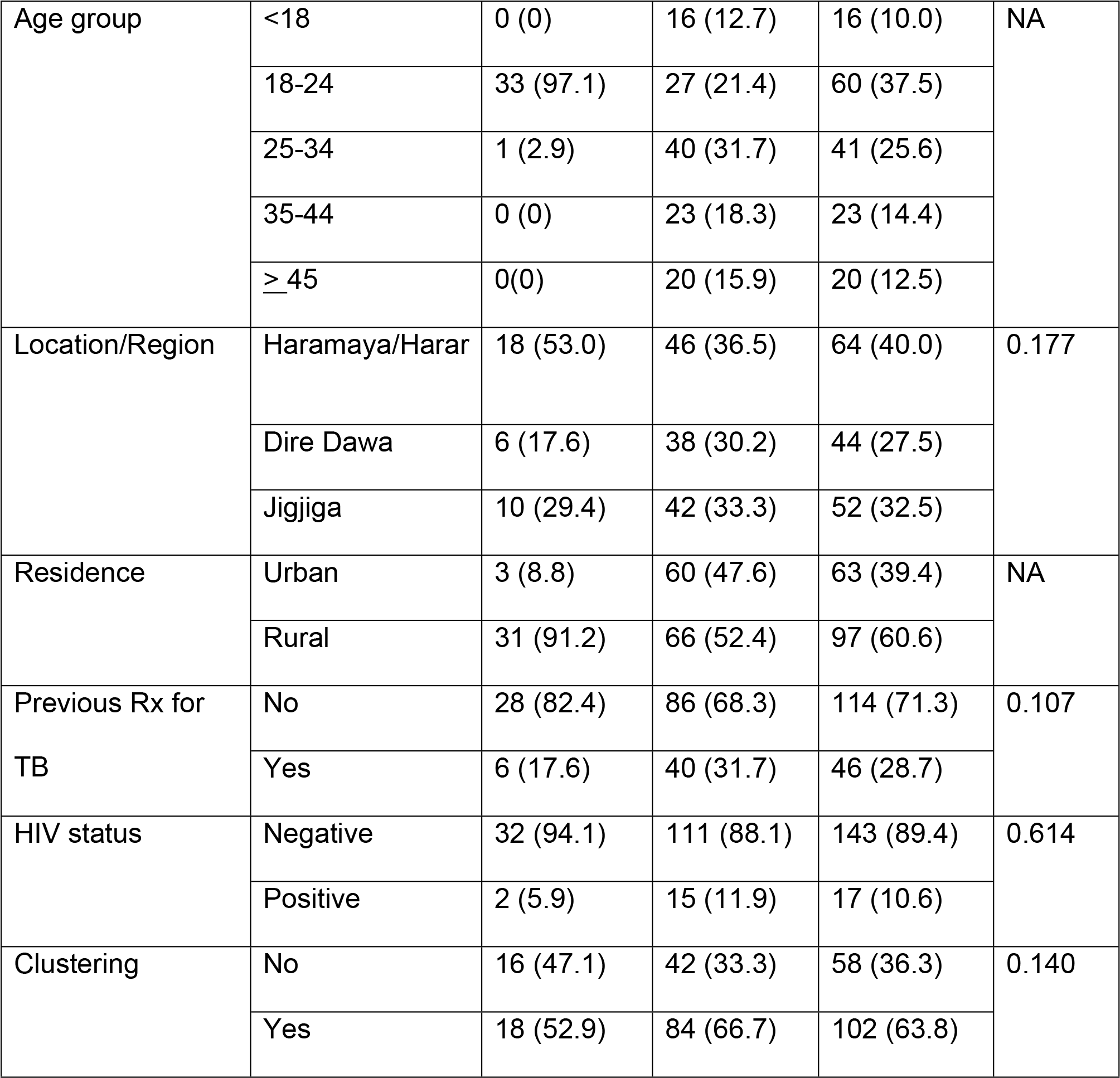

### Phylogenetic analysis of *M. tuberculosis* strains

The two predominant MTBC lineages in this study were lineage 4 (Euro-American, 71.3%) and lineage 3 (Delhi-CAS, 22.5%). MTBC strains classified as lineage 1 (East African Indian), and lineage 2 (Beijing) were only identified in six and four patients, respectively (Table 2).

**Table 2.**
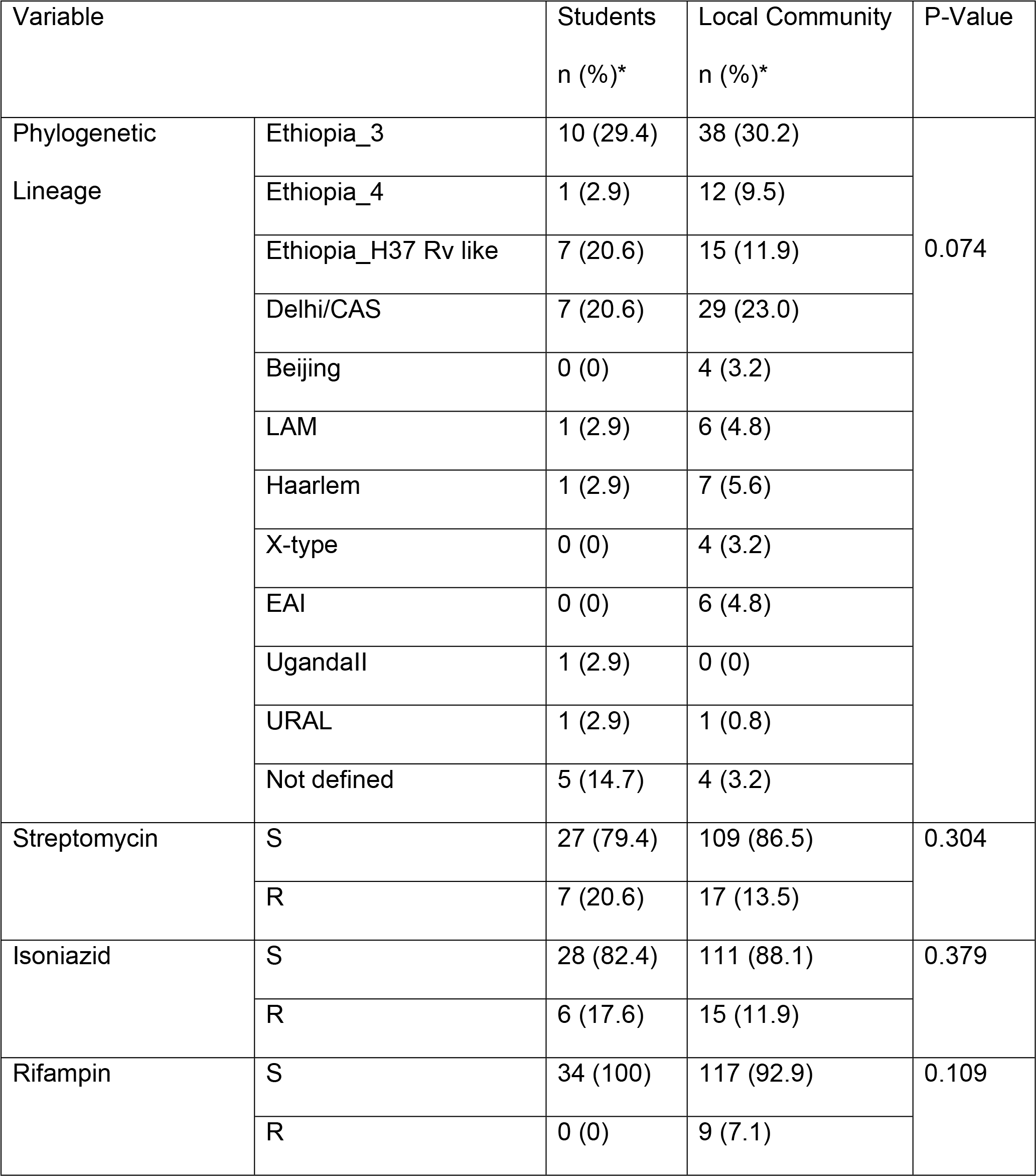
Phylogenetic Sub-lineages and Drug Sensitivity patterns of MTBC strains isolated by study participants, eastern Ethiopia, May 2016 to April 2017

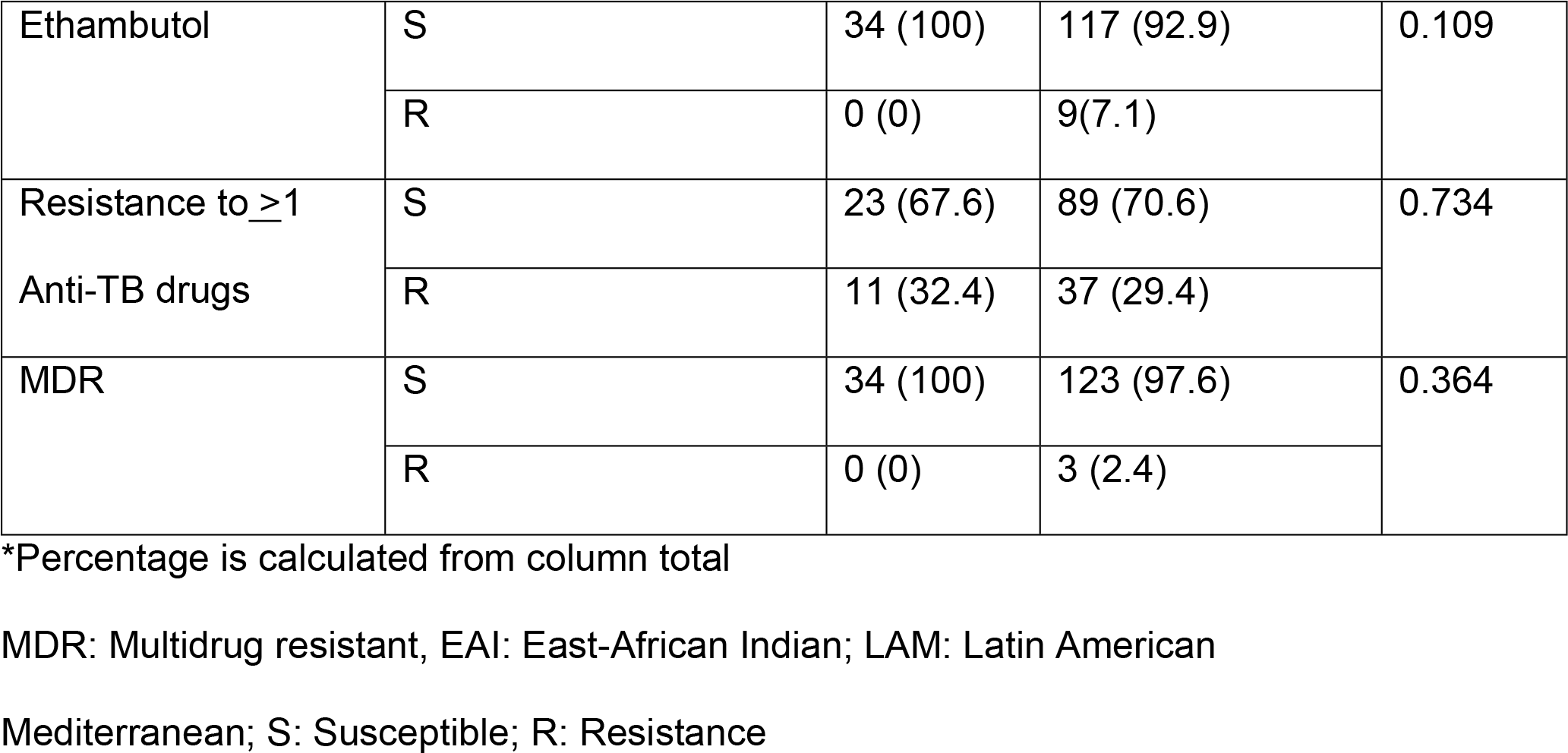

The most prevalent MTBC genotypes within the Euro-American super-lineage were Ethiopia_3 (29.4%) and Ethiopia-H37Rv-like (12.8%), both previously described among Ethiopian patients with PTB [10], and TB lymphadenitis [21]. Based on the phylogenetic structure/topology (UPGMA and MST-based), we further termed the third largest monophyletic lineage 4 group “Ethiopia_4” (8.1%) accordingly (Fig 1). Ethiopia_4 strains are closely related to Ethiopia_3 strains but with a distinct Spoligotyping pattern. Both, Ethiopia_3 and Ethiopia_4 strains, have a shared common ancestor with TUR genotype strains, but with unique Spoligotyping patterns, justifying their own nomenclature in the context of the molecular epidemiology in Ethiopia.

<u>Fig 1. Minimum spanning tree based on MIRU/VNTR profiles of 24- loci of 160 MTBC isolates in eastern Ethiopia. A circle representing a specific genotype is divided in to the number of strains clustering in it.</u>

MTBC strains classified as Beijing, East Africa India (EAI), or X-type were isolated exclusively from community members and not identified among university students. Particularly Beijing strains were only isolated from members of the surrounding Dire Dawa community. Nine MTBC isolates (5.6%) could not be classified within any of the known genotypes but these strains were part of lineage 4 with an unknown sub-group and are labeled “Not defined”.

### Molecular MTBC clusters and associated risk factors

The overall cluster rate of MTBC strains derived from all patients was 63.8%, including 21 clusters with 2 to 22 patients. Twelve clusters contained at least one university student. The overall Recent Transmission Index (RTI) was 50.6% (Table 3). In a univariate regression model, we found the following demographic and treatment related factors associated to clustered cases: female gender (P=0.004), urban residence (P=0.012) and those without prior TB diagnosis of active TB disease (P=0.001). Additionally, MTBC strains classified as Ethiopia_3 were significantly more abundant among clustered cases compared to other MTBC genotypes/lineages (Table 4, Fig 2).

**Fig. 2.**
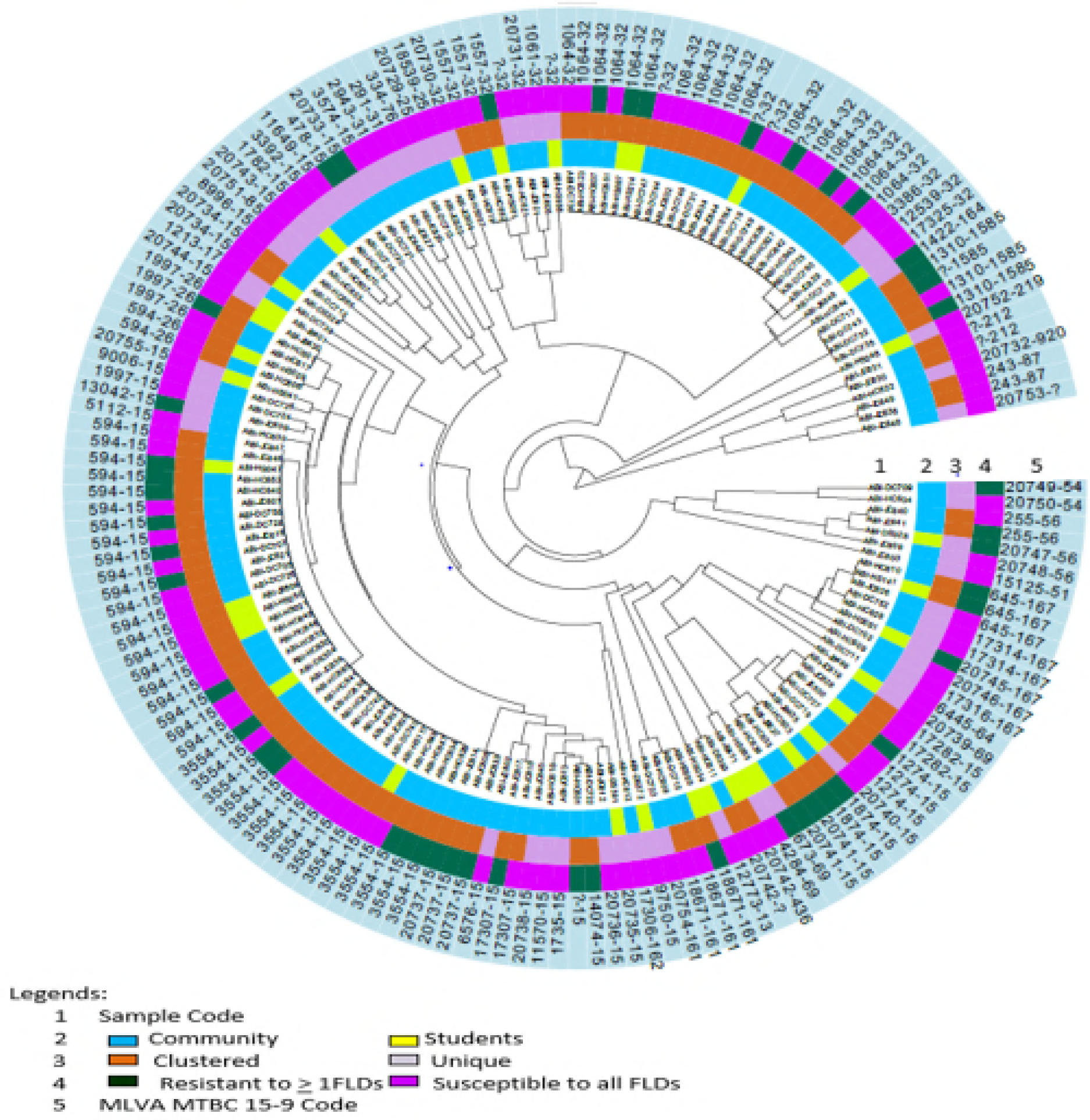
Radial UPGMA tree based on MIRU/VNTR 24-loci copy number. patient origin (Community or student), Clustering, Resistance to one or more FLDs (First line Anti TB drugs), and MLVA MTBC 15-9 Code.

**Table 3.**
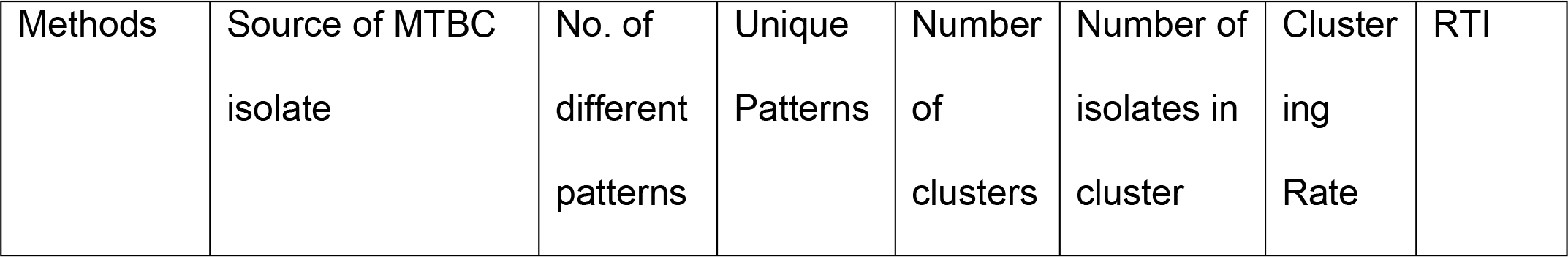
Clustering rate and Recent Transmission Index (RTI) analysis using Spoligotyping and MIRU-VNTR 24-loci methods for local community, university students and all study participants, eastern Ethiopia, May 2016 to April 2017

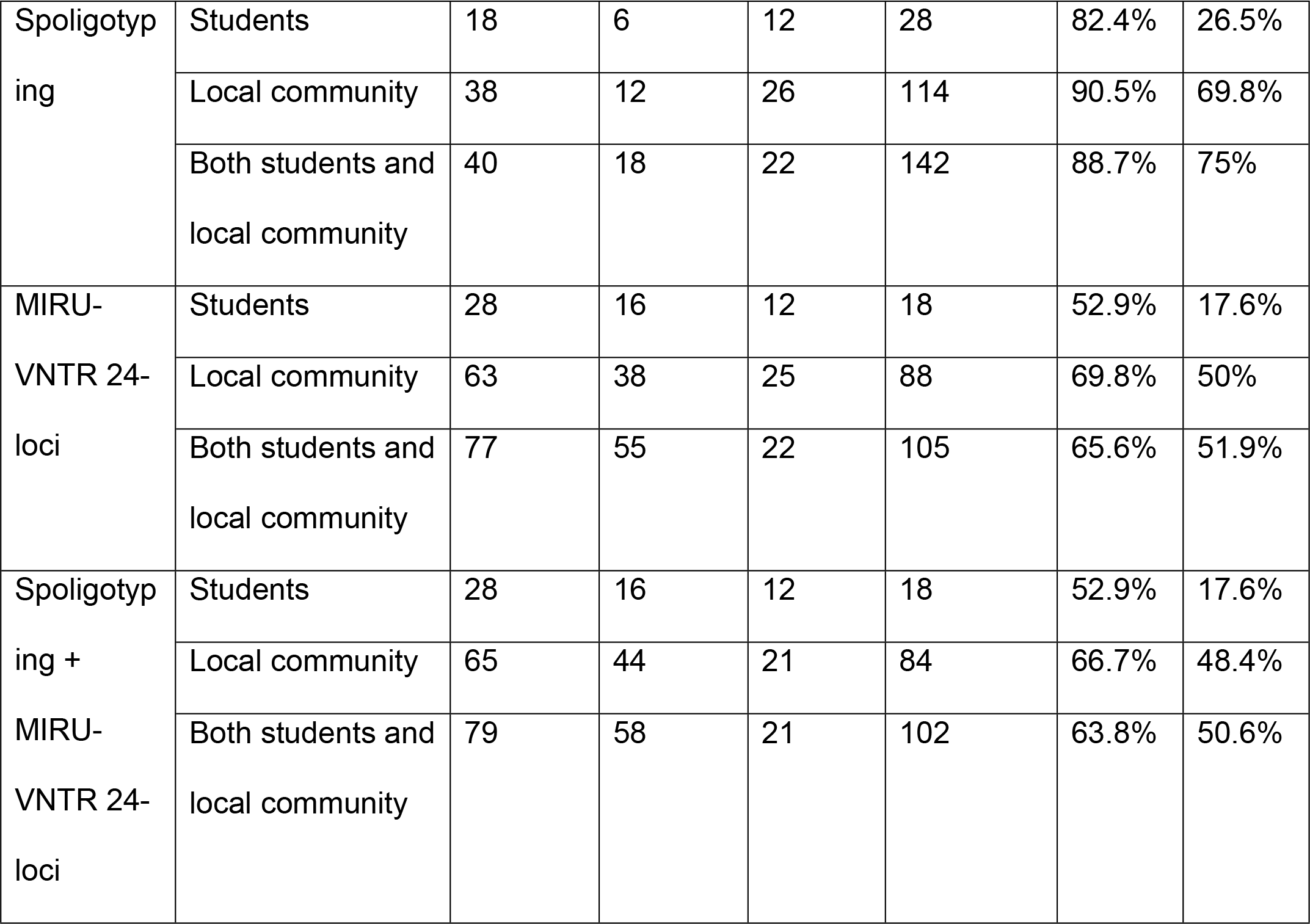

<u>Fig 2. Radial UPGMA tree based on MIRU/VNTR 24-loci copy number.</u>

**Table 4.**
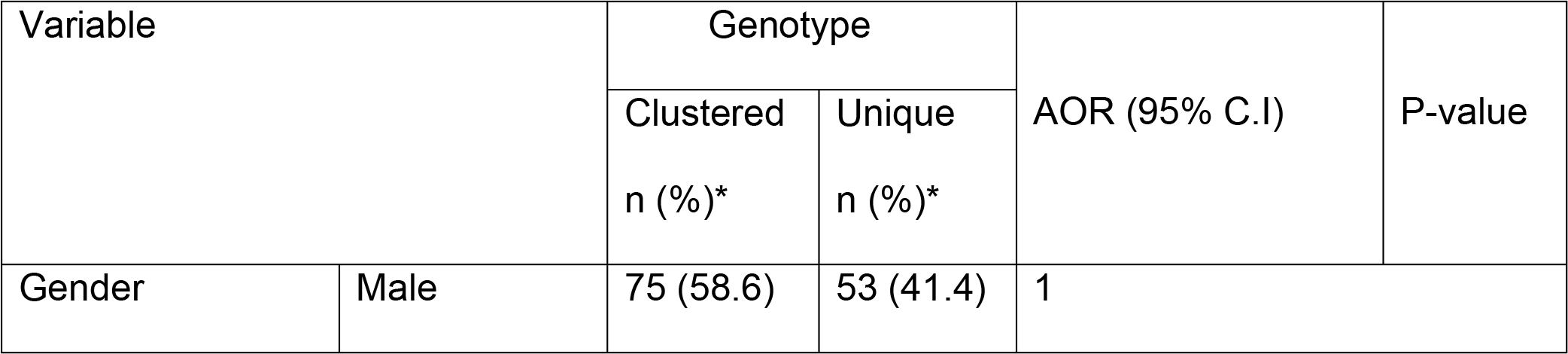
Factors associated with MTBC strain clustering, eastern Ethiopia, May 2016 to April 2017

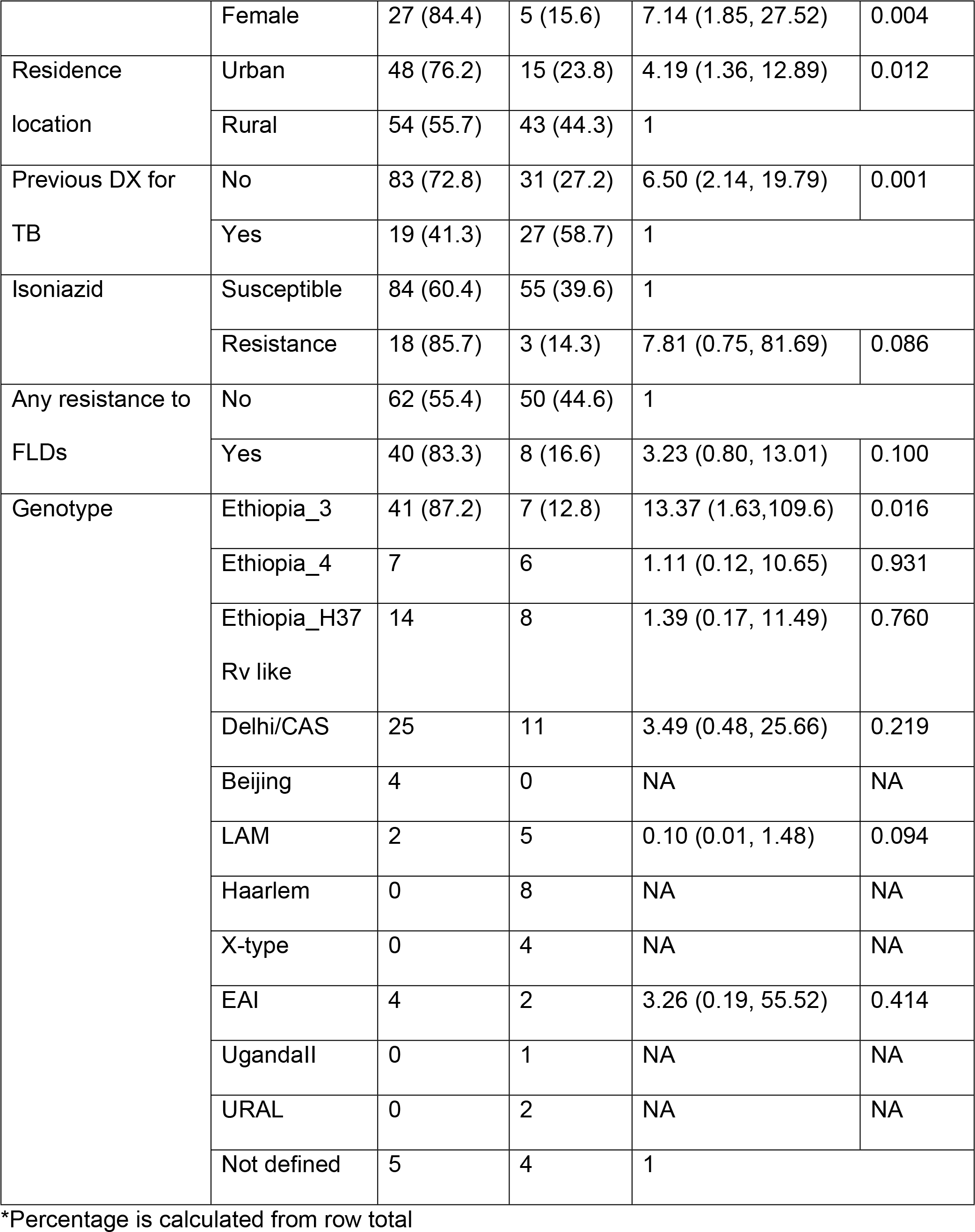

There was no a statistically significant difference in the proportion of clustered strains between university students and local community (p=0.142) (Table 1). With regard to the university cohort, 18/34 (52.9%) patients were part of a molecular cluster. 17 (94%) lived in an area at least 400 km away prior to attending university. There were two clusters with multiple student cases. One cluster contained three students, all of whom were attending Haramaya University. Two of the three students shared a common area of study, but none were living in the same dormitory or building. The other cluster contained five student cases, with three of the students attending Haramaya University and the other two attending Jigjiga and Dire Dawa Universities, respectively. None of the students in this cluster shared a clear epidemiologic link. Of the eight students clustered with other student cases, six (75%) reported recent exposure to someone with a cough, but none had a known exposure to an active TB case on campus. The RTI among students was 17.6% (Table 3).

### Patterns of *M. tuberculosis* drug resistance

Forty-eight (30%) of the study participants were resistant to one or more first line anti TB drugs. Among the first line anti-TB drugs examined, resistance to streptomycin and isoniazid was most common at 15%, and 13.1%, respectively. Rates of resistance to at least one first line anti-TB drug were similar in students and the community (32.4% vs 29.4%, p=0.734) (Table 2). Few participants had multidrug resistant (MDR) TB, with three cases occurring in community members (infected with Ethiopia_3, Ethiopia H37 Rv like, and LAM strain, respectively) and none occurring among university students.
Ethiopia_3 strains were observed with a higher proportion (9.4%) of resistance to at least one first-line drug compared to other MTBC genotypes (Table 5).

**Table 5.**
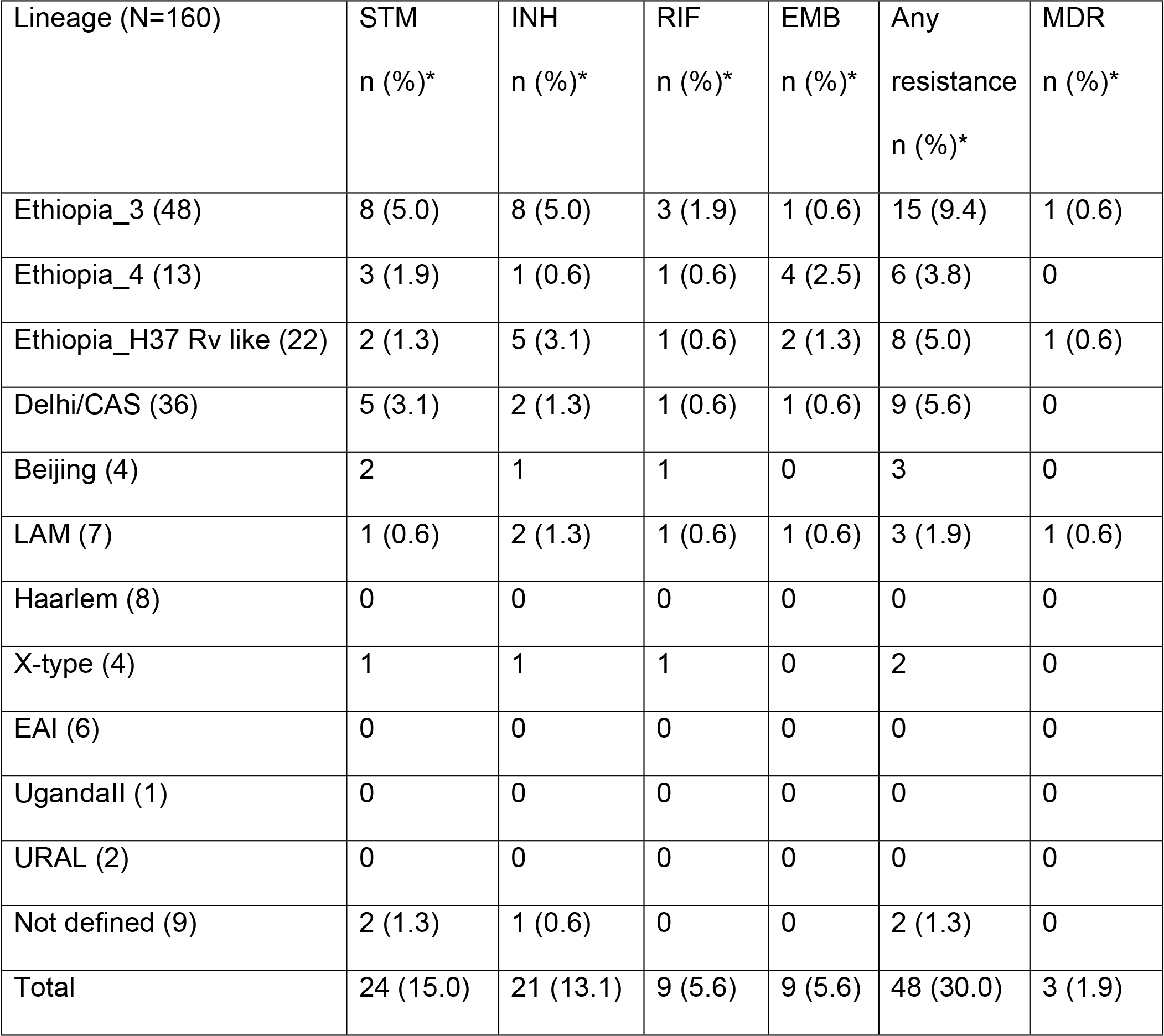
Patterns of drug resistance to first line anti-TB drugs by MTBC lineage eastern Ethiopia, May 2016 to April 2017

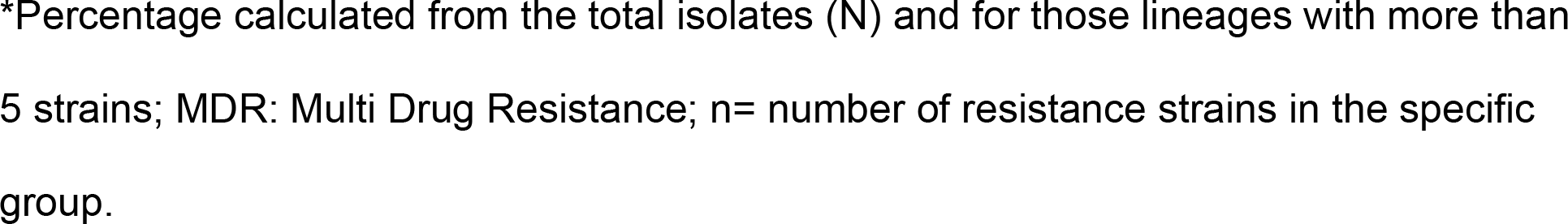

## Discussion

By using molecular MTBC strain typing (24-loci MIRU-VNTR typing and Spoligotyping) in combination with epidemiological investigations we point out the importance of MTBC Ethiopia_3 strains in the context of recent transmission and drug resistance in Eastern Ethiopia. Cluster rates among students and other community members were similar (52.9% and 66.7%, respectively [P =0.142]) and all clusters with university cases had at least one case from the community cohort associated. Being aware of the limitations mediated by the small sample size and the cross-sectional sampling scheme, our data suggest that MTBC transmission is still a major public health concern in Eastern Ethiopia and is not limited to congregate living facilities, e.g. universities or prisons [22].

Most university students with active TB originated from areas at least 400 km from the university campuses which might well be the source of the MTCB infection of non-clustered cases. Given the high cluster rate and the links to TB cases in the local community also suggests a significant proportion of active TB disease among students is due to transmission either on campus or in surrounding areas. Students within a molecular cluster were not observed with a direct epidemiologic link, indicating rather random short-term exposures as source of infection. These findings indicate that TB transmission is not within university campuses but is rather a community associated phenomenon. Larger scale molecular surveillance of MTBC strains in the region may be required to comprehensively characterize community transmission dynamics and design the most impactful interventions to interrupt transmission.

Molecular clusters defined by the applied genotyping methods are a surrogate marker for recent transmission and epidemiological linked cases [23]. The high clustering rate of 63.8% in this study indeed indicates that the majority of active TB cases in Eastern Ethiopia are due to recent transmission and not reactivation. Studies from other parts of the world, including South Tawara, Kiribat [24], have shown an even higher clustering rate of 75.3%. Although there were no previous studies conducted in eastern Ethiopia using the standard combined application of 24 loci MIRU/VNTR and Spoligotyping methods, a study from Northwestern Ethiopia demonstrated a clustering rate of 45.1% [10], differences might be explained by different living environments in the areas studied (e.g. rural vs. metropolitan setting) or other demographic differences (e.g. age, gender) [22].

Here, multivariable analysis demonstrated several demographic and clinical factors were associated with clustering. Females were about 7 times more likely than males to be a part of a cluster. This finding is similar to previous studies from Ethiopia [25] and Botswana [26] and could be linked to an increased tendency of females spending more time in close contact with their relatives in social sittings like market places compared to males. Urban residents were more than four times more likely part of a cluster, compared to their rural counterparts, which could be a result of the dense living conditions in cities. However, the burden of tuberculosis was higher among rural residents, which may reflect higher rates of poverty and less access to health care among the rural population.

Univariate analysis showed that strains with resistance to at least one FLD were more likely to be part of a cluster and were strongly associated with the Ethiopia_3 genotype. This is consistent with previous findings that even mono-resistant isolates may demonstrate selective pressure for MTBC transmission in Ethiopia [10, 27]. Similar to previous studies in Eastern Ethiopia [29], the prevalence of MDR TB was low. As recent data from South Africa suggests person-to-person transmission of drug resistant MTBC strains is the primary mechanism for the propagation of drug resistance, it will be important to improve public health programs to minimize transmission of these isolates [28].

The predominant MTBC lineages in this study were lineage 4 followed by lineage 3; findings that are similar to other studies conducted in Ethiopia [10, 19, 22]. We also demonstrate that local sub-linages such as the newly described “Ethiopia_4” and the closely related to Ethiopia_3 (both part of lineage 4 and related to TUR genotype strains) dominate among Ethiopian TB cases but do not play a major role in the global TB epidemic. This might be another example of a specialized, locally adapted MTBC strain type as recently suggested by Stucki and colleagues for the lineage 4 strains [30]. However, it is also important to note that strains with the Ethiopia H37Rv-like genotype can be found in many other world regions [24, 31, 32], and shares a common ancestor with the H37Rv laboratory reference strains (MTBC lineage 4.7, 4.8, and 4.9 according to Coll and his colleagues [33].

Strains from the Ethiopia_3 sub-lineage were more likely to be part of a cluster, indicating active transmission in the study area; an observation that was also found in Northwestern Ethiopia [10] and another study from Eastern Ethiopia [21]. The association between the Ethiopia_3 genotype and resistance to at least one first line drug in our study might be one factor that contribute to the expansion of this strain type. This finding highlights the need to conduct a large scale and more detailed characterization of the Ethiopia_3 sub-lineage in Ethiopia. Studies conducted in Northern, Northwestern and Southwestern Ethiopia have found the Dehli/CAS to be the predominant sub-lineage [10, 21, 27, 34], which may be attributable to these regions bordering Sudan, where Dehli/CAS was the most prevalent MTBC genotype [35]. All active TB cases caused by the Beijing strain, which is associated with high virulence, multidrug resistance and increased mortality [36], originated from the Dire Dawa community, which also supports the need to monitor the disease in the region. This study did not reveal any lineage-7 isolates (also referred to as “Ethiopia_1” in previous studies [10. 21]), which is in agreement with previous work demonstrating the predominance of lineage-7 in the northern part of the country [37].

This study is subject to several limitations. Our study in the local community included only smear positive PTB cases who have access to and visited the study hospital laboratories, which may not be representative of all active TB cases in the region. While our study is the first to provide insight about how TB is transmitted among persons attending university in eastern Ethiopia, the number of isolates from university students available for genotyping was small, limiting the ability to draw definitive conclusions about TB transmission.

## Conclusion

This study suggests TB-transmission in Ethiopia still is a major public health concern, and that transmission among college students is not limited to the university settings but also occurs in the general community. While the burden of MDR-TB in eastern Ethiopia remains low, we present evidence that the sub-lineage Ethiopia_3 is linked to recent TB transmission and also associated with resistance to at least one first-line drug. Country-wide comprehensive molecular surveillance and DST profiling of MTBC strains may be useful to guide ongoing and future TB control programs.

## Supporting Information

**S1 Fig.** Neighbor joining (NJ) phylogenetic tree based on 24 loci MIRU-VNTR profiles of 160 MTBC isolates from eastern Ethiopia (PDF) in relation to the MTBC reference collection hosted on miru-vntrplus.org.

## Acknowledgements

We would like to express our appreciation to the administrations of the respective universities and hospitals where the study was conducted. We thank all study participants and data collectors of the study. We are grateful to Harari Health Research and Regional Laboratory and Ethiopian Public Health Institute for their lab facility support. We acknowledge Anja Lüdemann, Tanja Struve-Sonnenschein, and Tanja Ubben for their support in MIRU-VNTR typing and Spoligotyping. We appreciate all-rounded support from department of Microbial, Cellular and Molecular Biology, Addis Ababa University.

## Authors contribution

AM conceived and designed the study. AM, BP, JC, GA and AA reviewed the proposal, contributed to the experiment, analysis and interpretation of the data, as well as the composition of this manuscript. AM, MM and SN participated in cluster analysis, strain classification, interpretation of the results and reviewed the initial and final manuscript. AM and DA participated in Drug Sensitivity Testing, interpretation of results and reviewed the manuscript. All authors read and approved the final manuscript.

